# Genome-wide Analysis of the VQ Gene Family and Expression Profiling in *Sorghum bicolor*

**DOI:** 10.64898/2026.11.05.764736

**Authors:** Fuwei Wang, Wei Zhang, Fangquan Zhang, Lei Wu, Qiushuo Wang, Mingyue Wang

**Affiliations:** Shandong Hi-speed Urban and Rural Construction Development CO., Ltd., Jinan, 250014

**Keywords:** Salt stress, VQ gene, Sorghum, Evolutionary analysis, Protein-protein interaction

## Abstract

Soil salinization severely restricts agricultural productivity, making the stress-tolerant sorghum an ideal model for studying crop adaptation. VQ motif-containing proteins are plant-specific transcriptional co-regulators vital for stress responses, yet their roles in sorghum remain underexplored. Here, we identified 35 *VQ* genes in sorghum, classifying them into seven subfamilies. Evolutionary analysis indicated that the family’s expansion was primarily driven by segmental duplication. Notably, synteny analysis revealed a lack of collinearity between sorghum and Arabidopsis *VQ* genes, suggesting significant functional divergence during monocot evolution. Expression profiling showed distinct patterns among VQ members, with *SbVQ34, SbVQ39* adn *SbVQ40* significantly upregulated under salt stress. Furthermore, protein-protein interaction analysis highlighted SbVQ39 as a potential key regulator in sorghum salt tolerance. This study provides novel insights into sorghum’s salinity tolerance mechanisms and establishes a foundation for utilizing *VQ* genes in breeding salt-tolerant crops.

## Introduction

Saline-alkali land is one of the important land resources in the world. Its soil salinity is high, its land use value is low, and it is not conducive to the growth and development of crops. Therefore, the utilization of saline-alkali land has always been an important research topic in the agricultural field. As a traditional dominant crop in my country, sorghum has strong environmental adaptability and excellent quality, and has received more and more attention in recent years. Sorghum (*Sorghum bicolor* L.) is a drought-tolerant and climate-adaptive crop that originated more than 60 centuries ago and was first domesticated in northeastern Africa^[1]^. It is an important cereal crop and is the fifth most cultivated cereal in the world, after barley, corn, wheat and rice ^[2]^.Sorghum has strong drought tolerance and salt-alkali tolerance, and is widely planted in arid-semi-arid areas and high-saline-alkali areas in Asia, Africa, and North America. Under the same water conditions, sorghum has higher water use efficiency than other crops such as corn, which makes it a major food source in the world^[3,4]^.

Valine-glutamine (VQ) protein is widely present in plants, and VQ members have been identified in Arabidopsis, rice, corn, lychee, potato and other plants^[5-9]^. Valine–glutamine (VQ) motif-containing proteins are transcriptional regulators that interact with known transcription factors (TFs), such as WRKY, PIF, and bHLH-type TFs, that are involved in regulating plant growth^[10,11]^. This interaction depends on the highly conserved VQ motif (FxxhVQxhTG) in VQ proteins^[12]^. Among these amino acid residues, x is any amino acid, h is a hydrophobic residue, and the last three residues show a variety of differences; depending on those differences, the VQ motif can be divided into different types, namely LTG, FTG, VTG, YTG, ITG, ATG, LTS, LTD, LTR, LTV, and LTA^[12]^. Aside from the highly conserved VQ motif, the amino acid sequences at other positions of VQ proteins show differences, leading to functional diversity among VQ family members^[13]^. Studies have shown that most VQ proteins are located in the nucleus, and some exist in the cytoplasm or membrane system, suggesting that they may be involved in transcriptional regulation or signal transduction^[14-15]^.

VQ proteins are not only involved in plant growth and development, but also in regulating plant responses to biotic and abiotic stresses. An increasing number of studies have shown that VQ acts as a transcriptional regulator in a variety of biological processes and then interacts with various transcription factors. In Arabidopsis plants, the PeVQ28 protein interacts with WRKY83 to enhance its transcriptional regulation of downstream salt/ABA-responsive genes, thereby positively regulating plant salt tolerance through the ABA-dependent signaling pathway^[16]^. In wheat plants, TaVQ4-D protein interacts with MPK3/MPK6 and plays a role in drought stress as its phosphorylation substrate, thereby promoting antioxidant enzyme activity and the expression of stress-related genes, and improving plant drought tolerance^[17]^. In apple plants, the MdVQ17 protein interacts with MdWRKY100 and inhibits the transcriptional activation ability of MdWRKY100 on the downstream target genes MdAKT1-like and MdPER57, thereby destroying Na+/K+ homeostasis and inhibiting ROS clearance, negatively regulating the salt tolerance of the plant^[18]^. In Arabidopsis thaliana plants, the StVQ31 protein regulates the expression of salt stress response-related genes including ERD, LEA4-5, At2g38905 and AtNCED3, enhancing the osmotic regulation ability and antioxidant homeostasis of cells, thereby improving the salt tolerance of transgenic plants^[19]^. In apple plants, the MdVQ8 protein interacts with MdWRKY75 and MdTCP46, enhancing the transcriptional activation ability of MdWRKY75 on MdMYB1 and promoting the MdTCP46-MdMYB1 interaction, thus promoting anthocyanin biosynthesis through dual pathways and improving the adaptability of plants under salt stress^[20]^. In Arabidopsis plants, overexpression of VQ1 protein enhances the plant’s multiple resistance to salt stress and pathogen infection by regulating the levels of abscisic acid and salicylic acid and differentially regulating a series of stress-related genes (such as up-regulation of MPK6, NPR1, PDF1.2, down-regulation of WRKY70, ABI1, etc.)^[21]^.

Although to date, there are extremely limited clues about how VQ is involved in the internal regulation of sorghum under salt stress, these studies will also prompt us to investigate the role of VQ in relation to salt stress in sorghum.

## Methods

### Identification of *VQ* gene family in sorghum

The identification of the *VQ* gene family in sorghum commenced with the retrieval of genomic and proteomic data for *sorghum bicolor* and *Arabidopsis thaliana* from the EnsemblPlants database^[22]^. An initial homology search was performed using the Blastp module and Advanced HMMER Search within TBtools-II^[23]^, utilizing Arabidopsis VQ protein sequences as queries against the sorghum protein database (E-value: 1e-5). To ensure accuracy, all candidate hits were rigorously filtered through conserved domain analysis to validate the existence of the signature VQ domain.

### Prediction of physical and chemical properties of sorghum VQ protein

Comprehensive bioinformatic analyses were conducted to elucidate the structural and functional attributes of the sorghum VQ protein family. Primary protein characteristics such as relative molecular mass, isoelectric point, and amino acid count were determined utilizing ExPASy online resources. Furthermore, tertiary structural data and functional domain architectures were retrieved and annotated from the UniProtKB/Swiss-Prot database.

### Systematic evolution, gene structure and protein motif analysis of sorghum *VQ* genes

A maximum-likelihood phylogenetic tree was constructed using TBtools-II to analyze the evolutionary divergence of VQ proteins in Arabidopsis and sorghum. The analysis employed 5,000 bootstrap replicates to assess node support. Subsequently, the tree topology was refined using ChiPlot. To characterize the protein architecture, conserved motifs were predicted using the MEME online server. Furthermore, the structural organization of sorghum *VQ* genes, including exon-intron boundaries and UTR locations, was annotated using the genome assembly from EnsemblPlants and graphically represented using GSDS. Collinearity analysis of *VQ* genes was performed using MCScanX, and the results were visualized using TBtools-II.

### Analysis of *cis*-acting elements in the promoter of sorghum *VQ* gene

The 2000 bp genomic regions upstream of the transcription start sites of *VQ* genes were submitted to the PlantCARE database to predict putative *cis*-acting regulatory elements. The identified stress-responsive and hormone-related elements were subsequently used to generate a heatmap illustrating the distribution and abundance of these motifs across the gene family.

### Expression analysis of sorghum *VQ* genes

Transcriptome datasets (NCBI GEO: GSE145748, GSE98817, GSE140928, GSE30249) and published RNA-seq data^[24]^ were retrieved to investigate *SbVQ* gene expression across various tissues and under abiotic stress. The expression profiles were normalized, and heatmap visualization was performed to illustrate the expression patterns.

### Prediction of Interactions between VQ and WRKY proteins in sorghum

All sorghum WRKY protein sequences, as detailed in Supplementary Table 1, were retrieved from previous studies^[25].^ The STRING database was employed to predict the interactions between VQ and WRKY family members, and the resulting interaction network was plotted using the Chiplot tool.

## Result

### Identification, protein characteristics, and gene structure analysis of the VQ gene family

Through homologous sequence alignment and conserved domain sequence analysis (Fig S1), we identified 40 *VQ* genes distributed across 10 sorghum chromosomes (Supplementary Table 1). Chromosome 1 harbored the highest number of genes (11), whereas chromosomes 5 contained only one each. The encoded proteins ranged from 84 to 453 amino acids in length, with molecular weights varying from 8.91 to 44.27 kDa. This variation in amino acid sequence contributes to diverse tertiary structures, which are crucial for VQ protein function. The isoelectric points (pI) ranged from 5.27 to 11.33, with the majority being alkaline, indicating an abundance of basic amino acids. Most VQ proteins exhibited instability indices greater than 40, suggesting the presence of destabilizing dipeptides. Notably, SbVQ12 and SbVQ28 displayed high aliphatic indices, indicating superior thermostability. Furthermore, all VQ proteins had negative GRAVY values, classifying them as hydrophilic. Specifically, SbVQ3, SbVQ4, SbVQ11, and SbVQ18 showed strong hydrophilicity, which is consistent with their putative nuclear localization as transcriptional regulators.

Motif analysis revealed that all identified proteins contain Motif 1, which corresponds to the conserved VQ domain (Fig. 1a-b). However, the scarcity of other shared motifs among VQ proteins indicates substantial sequence divergence outside this conserved domain. Even within the highly conserved and single short FxxhVQxhTG (h: hydrophobic amino acid, x: any amino acid) amino acid sequence motif^[26]^, considerable variation exists in the flanking regions (Fig. S2), suggesting potential functional diversification during evolution. Gene structure analysis showed that the majority of *VQ* genes are intronless, with only five genes containing introns of varying lengths. Notably, the relatively long introns in *SbVQ11* and *SbVQ20* may be associated with specific gene expression patterns, alternative splicing, and evolutionary selection (Fig. 1c).

**Figure 1.**
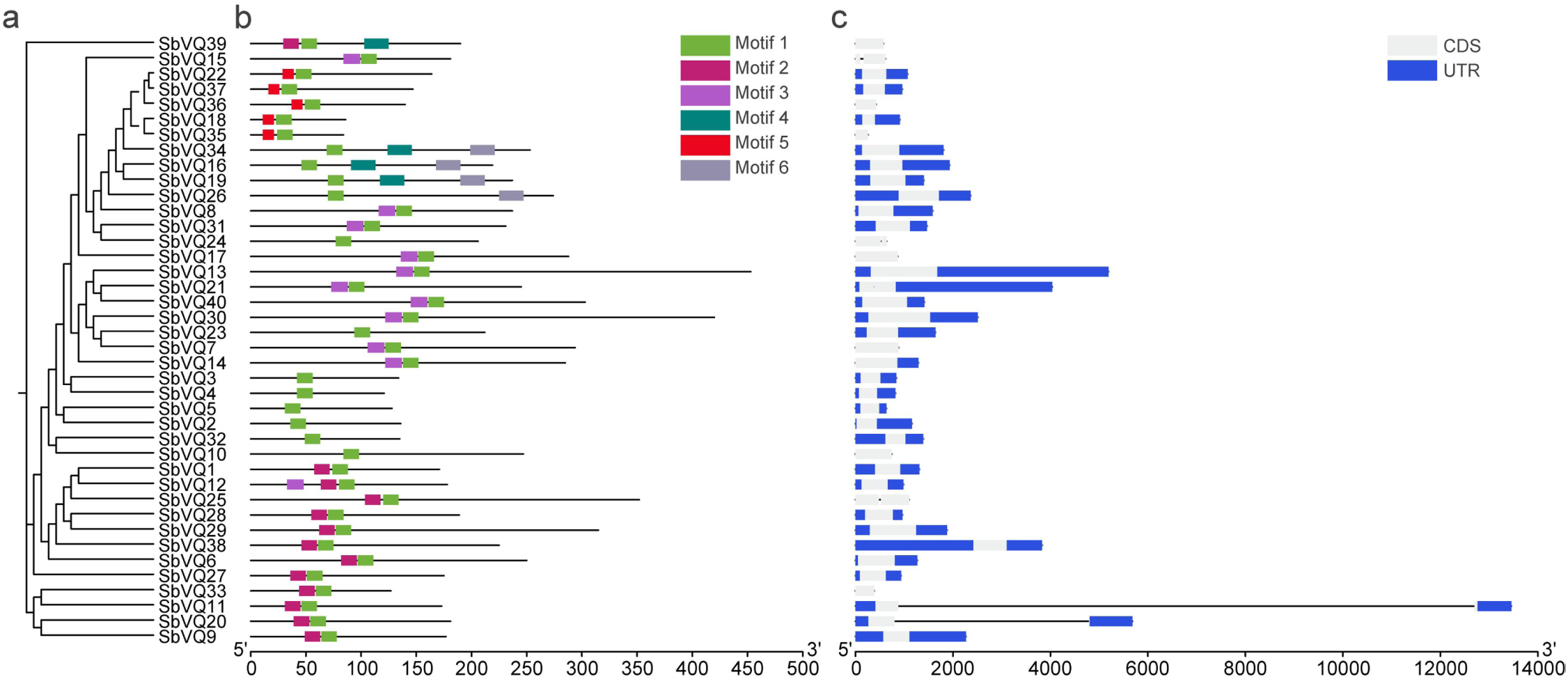
Analysis of motifs, and gene structures of the VQ family in sorghum. (a) Phylogenetic tree of VQ proteins; (b) Motif analysis of VQ proteins; (c) Gene structure analysis of *VQ* genes.

### Phylogeny, gene duplication, and synteny analysis of sorghum *VQ* genes

To investigate the evolutionary relationships of *VQ* genes in sorghum, we constructed a phylogenetic tree using VQ proteins from both *sorghum bicolor* and *Arabidopsis thaliana*. The sequences of VQ members from sorghum and Arabidopsis are listed in Supplementary Table 1. Based on the phylogenetic analysis, the sorghum *VQ* genes were classified into seven distinct subfamilies (Fig. 2). Among these, subfamily III contains the fewest members, with only three sorghum *VQ* genes, whereas subfamily IV is the largest, comprising eight genes. This evolutionary divergence suggests that the diversification of *VQ* genes may be closely associated with the adaptation of sorghum to complex environmental conditions.

**Figure 2.**
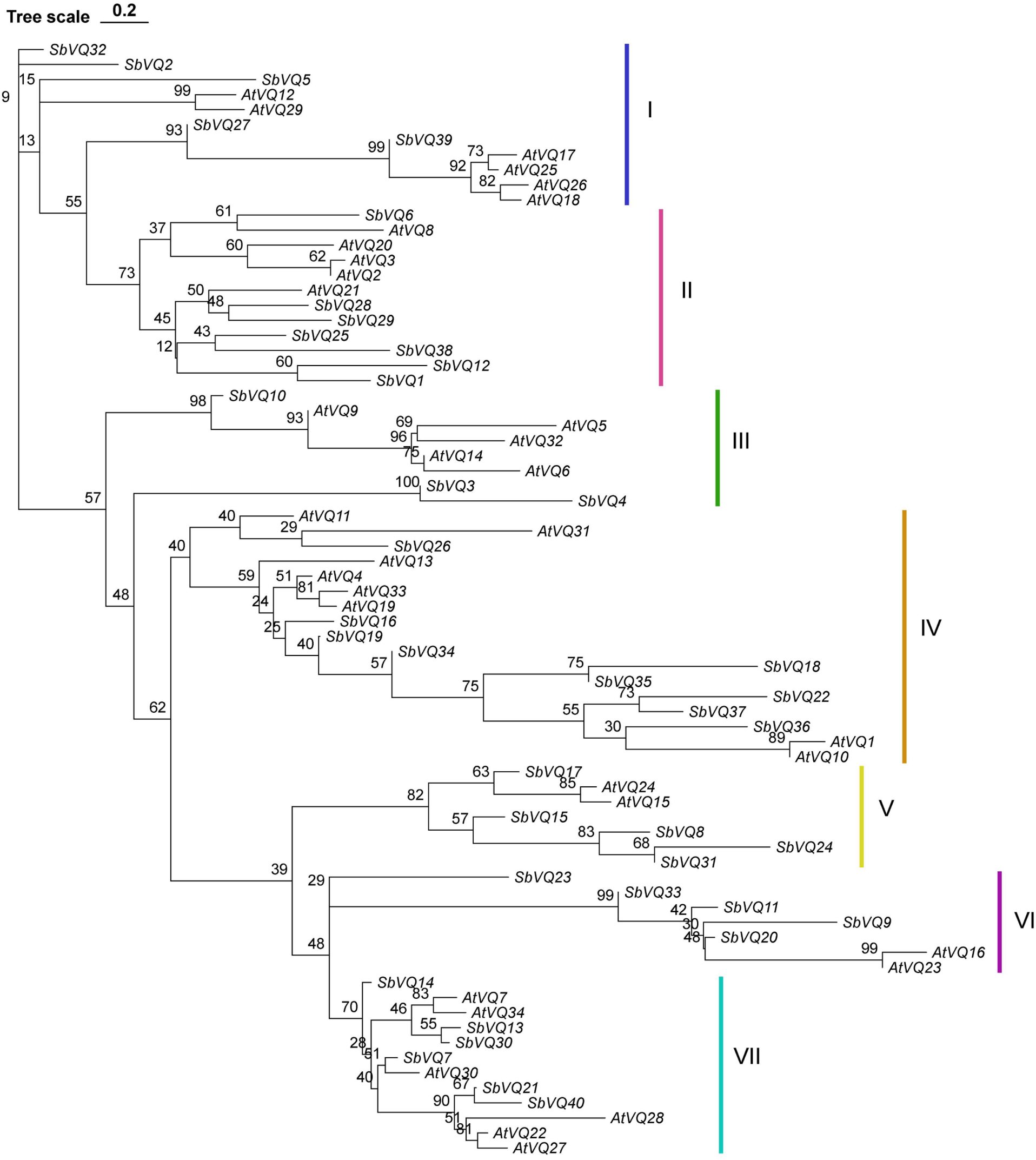
Phylogenetic relationships of VQ proteins between sorghum and Arabidopsis. The *VQ* genes were classified into seven distinct subfamilies. Different colored blocks represent different subfamilies.

Furthermore, we investigated the duplication patterns of the sorghum *VQ* genes. The analysis revealed a substantial number of segmental duplication events, suggesting that segmental duplication has been the primary driving force behind the expansion of the *VQ* gene family in sorghum. This increase in *VQ* gene copy number may have enhanced the capacity of sorghum to adapt to complex environmental conditions (Fig. 3a). Additionally, we performed a synteny analysis between sorghum, Arabidopsis, and maize. The results showed that only one syntenic gene pair was identified between sorghum and Arabidopsis, whereas a large number of syntenic genes were found between sorghum and maize. This highlights the close evolutionary relationship between sorghum and maize, and implies that *VQ* genes may have undergone significant functional divergence between monocots and dicots (Fig. 3b).

**Figure 3.**
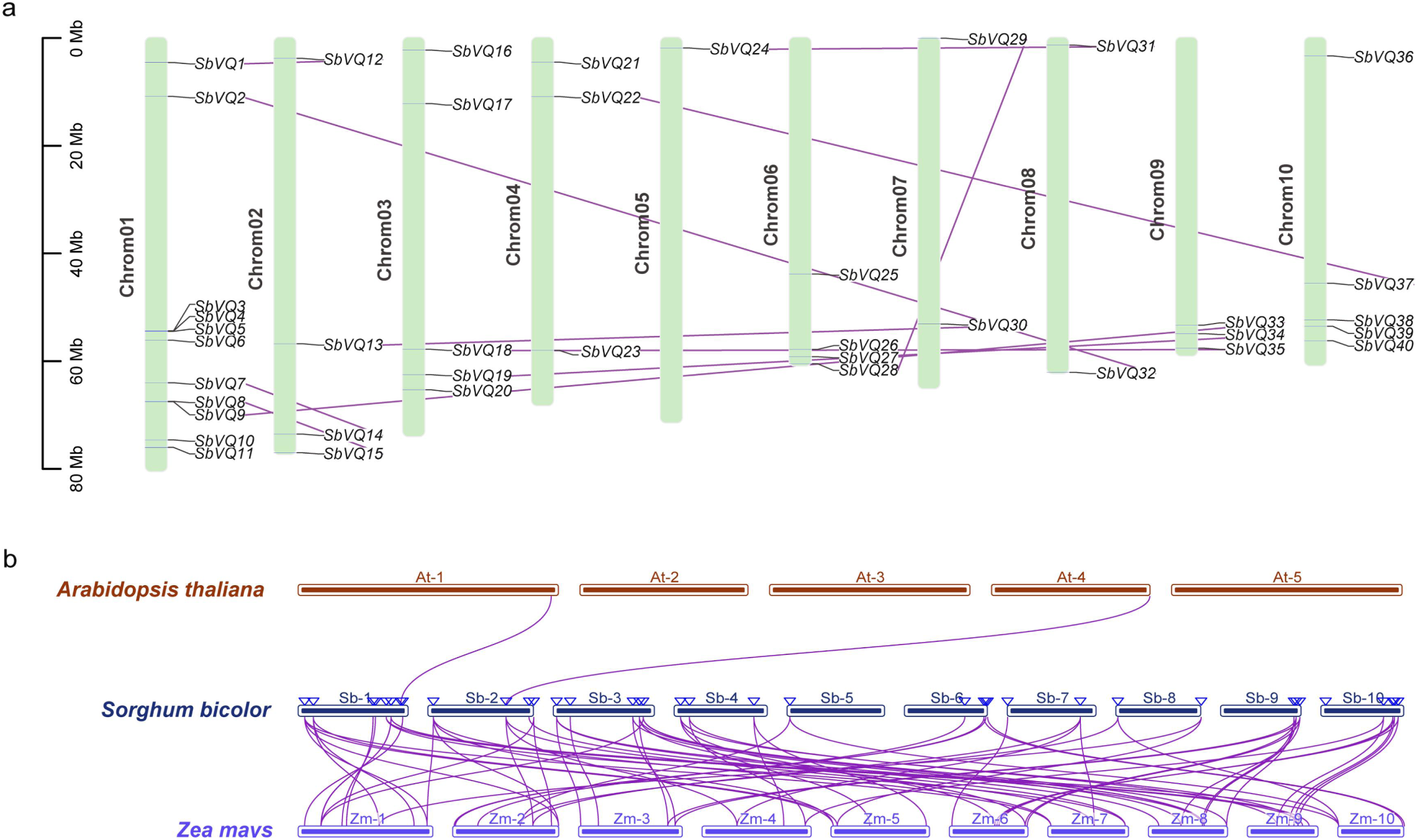
Syntenic analyses of *VQ* genes in sorghum. (a) Duplication event analysis of the sorghum *VQ* genes, highlighting the role of segmental duplication in gene family expansion. (b) Syntenic relationships of *VQ* genes among sorghum, Arabidopsis, and maize. Gray lines indicate syntenic blocks, while highlighted lines connect syntenic *VQ* gene pairs.

### Analysis of *cis*-acting elements in the promoters of sorghum VQ genes

To investigate the expression specificity and functional response mechanisms of *VQ* genes, we predicted and analyzed the *cis*-acting elements in the promoters of sorghum *VQ* genes. The identified *cis*-acting elements were categorized into three groups: hormone responsiveness, growth and development regulation, and biotic and abiotic stress responses. Regarding hormone-responsive elements, the TGACG-motif, which is associated with jasmonic acid (JA) signaling and stress responses, is widely present in the promoters of most *VQ* genes. The ABRE element, a binding site for bZIP transcription factors (e.g., AREB/ABF) involved in abscisic acid (ABA) signaling, is also highly prevalent. This indicates that the sorghum *VQ* family is primarily regulated by JA and ABA pathways. Notably, the abundance of ABRE elements in the *SbVQ26* promoter suggests its potential pivotal role in ABA signal transduction.

In terms of growth and development, G-box and MYB-responsive elements are the most prominent. The G-box is linked to photosynthesis and chloroplast development, and it can also synergize with ABA signaling to enhance stress tolerance. MYB-responsive elements are critical for secondary metabolism and defense. MYB transcription factors (especially the R2R3-MYB subfamily) regulate the phenylpropanoid pathway, driving the biosynthesis of flavonoids and anthocyanins. Additionally, they help coordinate responses to abiotic stresses like drought, salinity, and heavy metals via hormone signaling modulation. Specifically, the abundance of G-box elements in *SbVQ2/26* and MYB-binding sites in *SbVQ6/18/24/26* and *SbVQ35* implies their significant involvement in plant development and stress adaptation.

In terms of stress-responsive elements, as-1, MYC, and STRE motifs are the most prominent. The as-1 element shares its core sequence with the TGACG-motif, whereas the MYC binding site is targeted by bHLH transcription factors to mediate jasmonic acid signaling. Additionally, the highly conserved STRE plays a broad-spectrum role in stress responses. The abundance of these elements in *SbVQ6* implies its critical involvement in stress adaptation. Overall, promoter analysis offers key insights into gene regulation and expression features, laying a solid foundation for future functional studies (Fig. 4). Detailed information regarding promoter elements is presented in Supplementary Table 2.

**Figure 4.**
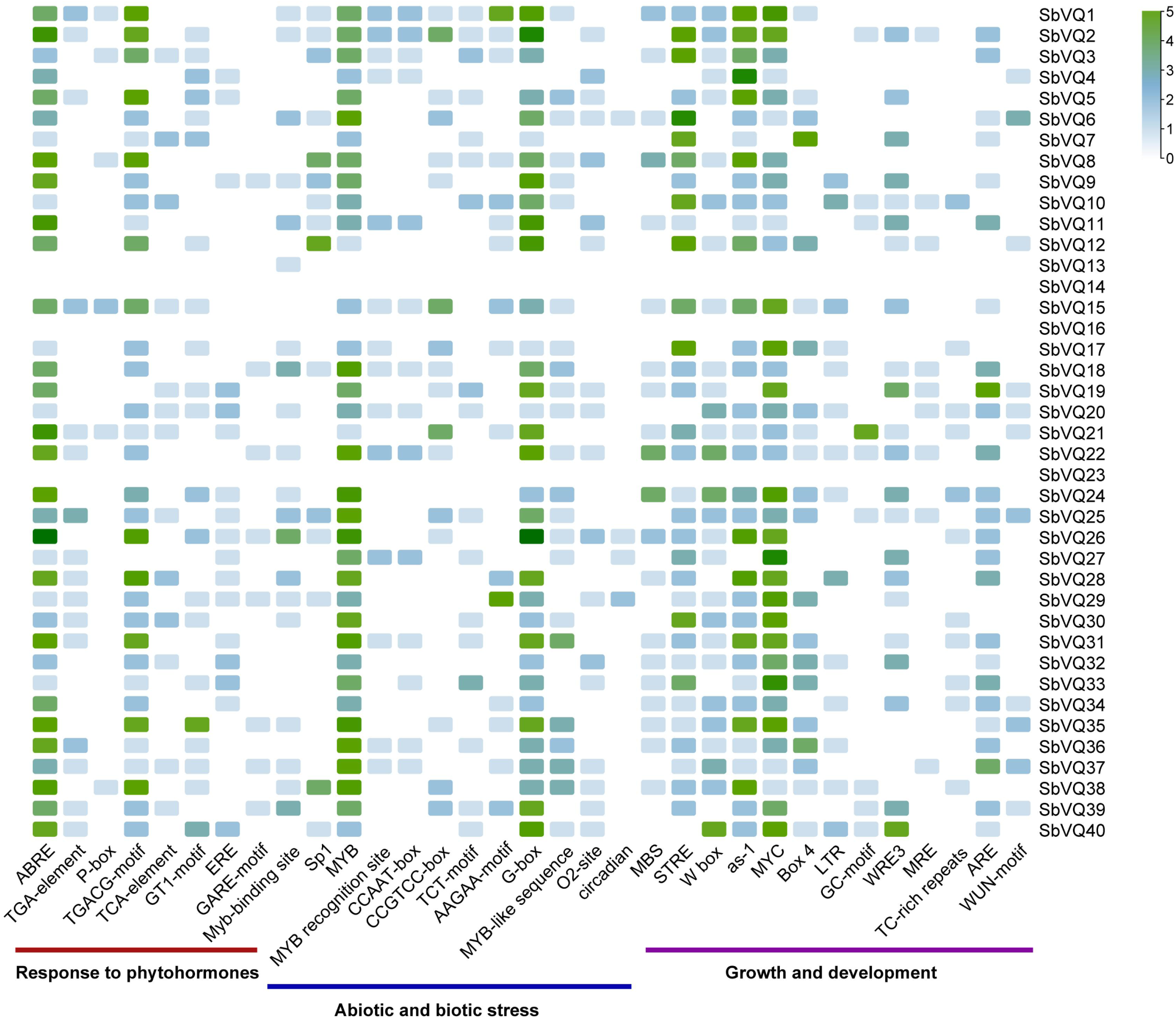
Analysis of *cis*-acting elements in the promoters of sorghum *VQ* genes. The elements are categorized into three groups based on their functional responses. The color intensity represents the number of elements, with darker colors indicating a higher quantity.

### Expression profiles of *VQ* genes across tissues, salt stress, and PEG treatment

To further investigate the functions of sorghum *VQ* genes, we analyzed their expression profiles using publicly available transcriptomic data. In the roots, *SbVQ9*, *SbVQ15*, *SbVQ21*, *SbVQ25*, *SbVQ32,* and *SbVQ40* exhibited relatively high expression levels. Notably, *SbVQ15* was induced by salt stress in the salt-tolerant line M-81E, while the expression of *SbVQ21 and SbVQ40* were significantly higher in the salt-tolerant line compared to the salt-sensitive line Roma. These findings suggest that both genes may be involved in the salt stress response of sorghum roots (Fig. 5a). In the stems, *SbVQ2*, *SbVQ19*, and *SbVQ34* showed elevated expression during the late stages of stem development, indicating their potential roles in stem maturation (Fig. 5b). In the leaves, *SbVQ40* displayed the highest expression level. Under severe salt stress (msq5), the expression of *SbVQ7*, *SbVQ10, SbVQ21*, *SbVQ34*, and *SbVQ39* was consistently upregulated, suggesting that these genes may be associated with the salt stress response in sorghum leaves (Fig. 5c).

**Figure 5.**
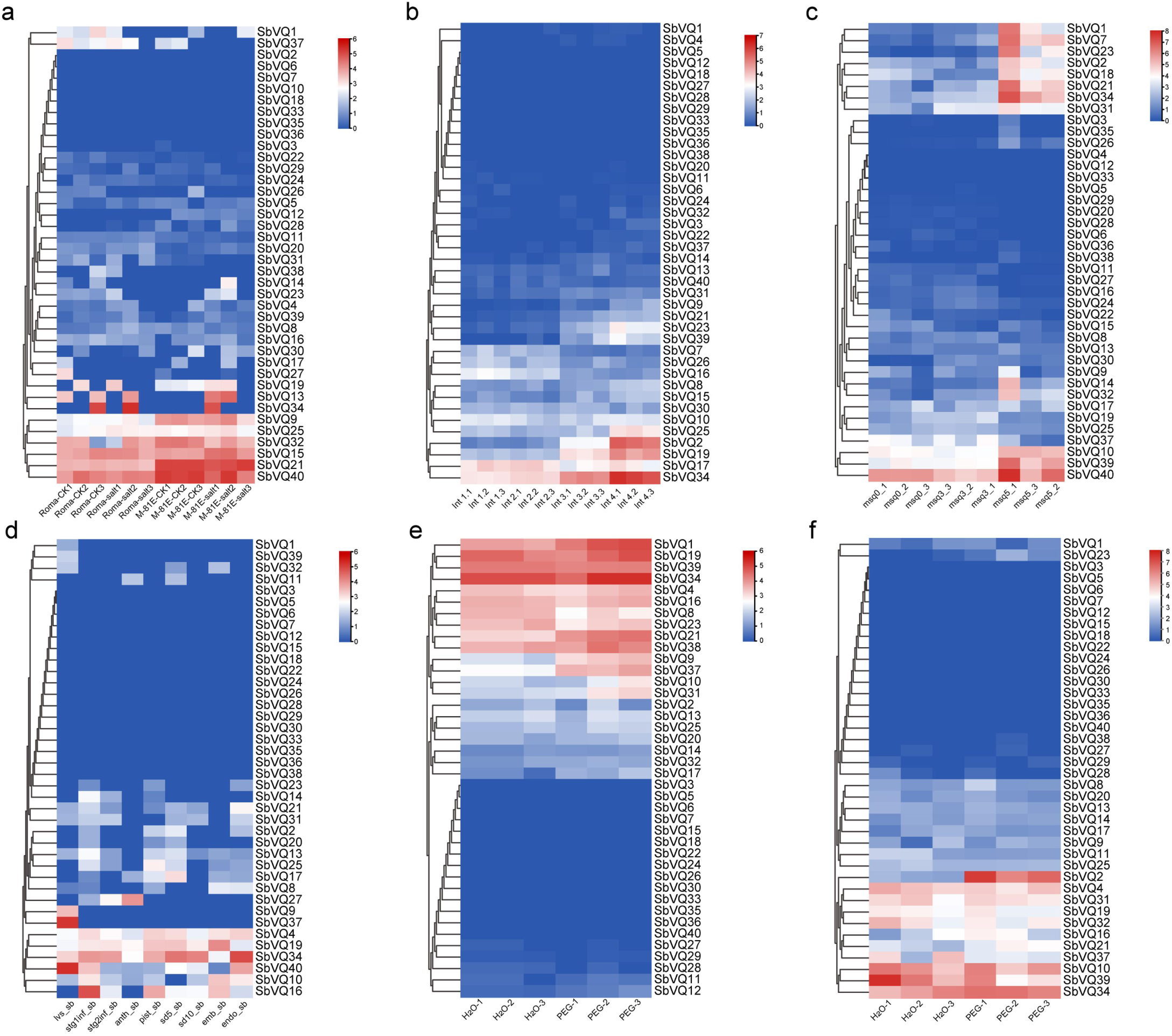
Expression pattern analysis of sorghum *VQ* genes. Color scale represents expression levels. Data are shown as log_2_(expression value) with columns clustered. (a) Expression profiles of *VQ* genes in the roots of the salt-tolerant sorghum line M-81E and the salt-sensitive line Roma. (b) Expression patterns of *VQ* genes at different developmental stages of sorghum stems. (c) Expression levels of *VQ* genes in sorghum leaves under mild and severe salt stress. (d) Expression patterns of *VQ* genes across various tissues and organs. (e) Expression profiles of *VQ* genes in sorghum roots under PEG treatment. (f) Expression profiles of *VQ* genes in the shoots of sorghum under PEG treatment.

Analysis of *VQ* gene expression across different tissues revealed that *SbVQ37* and *SbVQ40* exhibited significantly higher expression levels in leaves compared to other *VQ* family members. *SbVQ16* showed the highest expression in floral primordia, while *SbVQ34* was predominantly expressed in the endosperm. This tissue-specific expression pattern suggests that different *VQ* genes may play distinct roles in specific organs (Fig. 5d). Under PEG treatment, *VQ* genes in sorghum roots displayed diverse expression patterns: the expression of *SbVQ1*, *SbVQ9*, and *SbVQ37* were upregulated, whereas *SbVQ8* was significantly downregulated. In the shoot, *SbVQ2* and *SbVQ34* were upregulated, while *SbVQ37 and SbVQ39* were downregulated (Fig. 5e-f). These distinct transcriptional responses reflect the complex regulatory mechanisms of *VQ* genes in mediating drought stress. Overall, the comprehensive expression profiling provides valuable insights into the functional characteristics of individual members within the sorghum *VQ* gene family. The expression data of the sorghum *VQ* genes used in this study are listed in Supplementary Table 3.

### Interaction analysis between VQ and WRKY family members in sorghum

The tertiary structure of a protein dictates its function. Therefore, we first conducted a predictive analysis of the tertiary structures of representative VQ proteins from different subfamilies. The results revealed considerable structural variations among members of different subfamilies. Moreover, while the conserved VQ motif exhibited high confidence, the confidence levels for most other regions of the tertiary structures were relatively low. The conserved VQ motif is generally located at the center of the protein, forming an α-helix. This helical structure likely plays a pivotal role in the interaction between VQ proteins and other interacting partners (Fig. 6).

**Figure 6.**
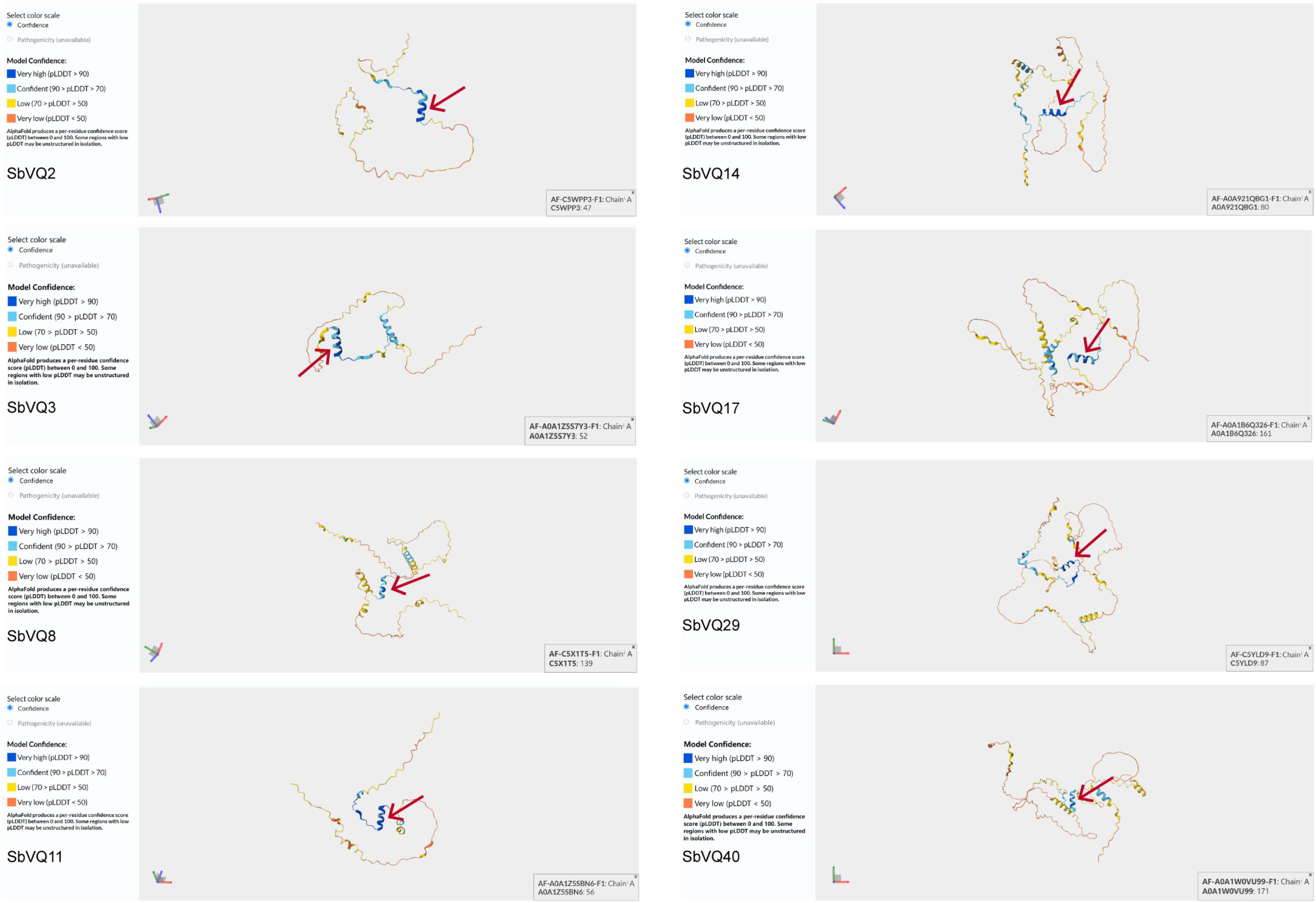
3D structures of representative VQ proteins from different subfamilies in sorghum. Red arrow points to the alpha-helix within the conserved domain of the VQ protein.

It is well established that most members of the *VQ* gene family interact with WRKY transcription factors. To preliminarily investigate the interactions between sorghum VQ and WRKY proteins, we predicted potential protein-protein interactions using a bioinformatics database. The predicted interactions are listed in Supplementary Table 4. The results suggest that SbWRKY51 (A0A1Z5RH15), SbWRKY52 (C5Y3H4), SbWRKY60 (A0A1B6PMC2), SbWRKY6 (A0A1B6QJ20), SbWRKY71 (A0A1B6PAY6), and SbWRKY42 (A0A1B6Q6V3) may serve as core genes within the sorghum WRKY family, while SbVQ10 (A0A1Z5SAZ3), SbVQ26 (A0A1Z5RGE7), SbVQ39 (C5Z655_SORBI), and SbVQ40 (A0A1W0VU99) could be core members of the sorghum VQ family. Furthermore, the WRKY and VQ families were found to interact through specific partners, including SbWRKY42 (A0A1B6Q6V3), SbWRKY66 (A0A1Z5RBX4), and SbWRKY79 (A0A1B6P7P6) (Fig. 7). However, we also noted that the interactions between a large number of *WRKY* genes and certain *VQ* genes were not predicted. Therefore, these computational predictions warrant further experimental validation. The sequences of the sorghum WRKY proteins used in this study are listed in Supplementary Table 1, and the interaction relationships between sorghum VQ and WRKY proteins are provided in Supplementary Table 4.

**Figure 7.**
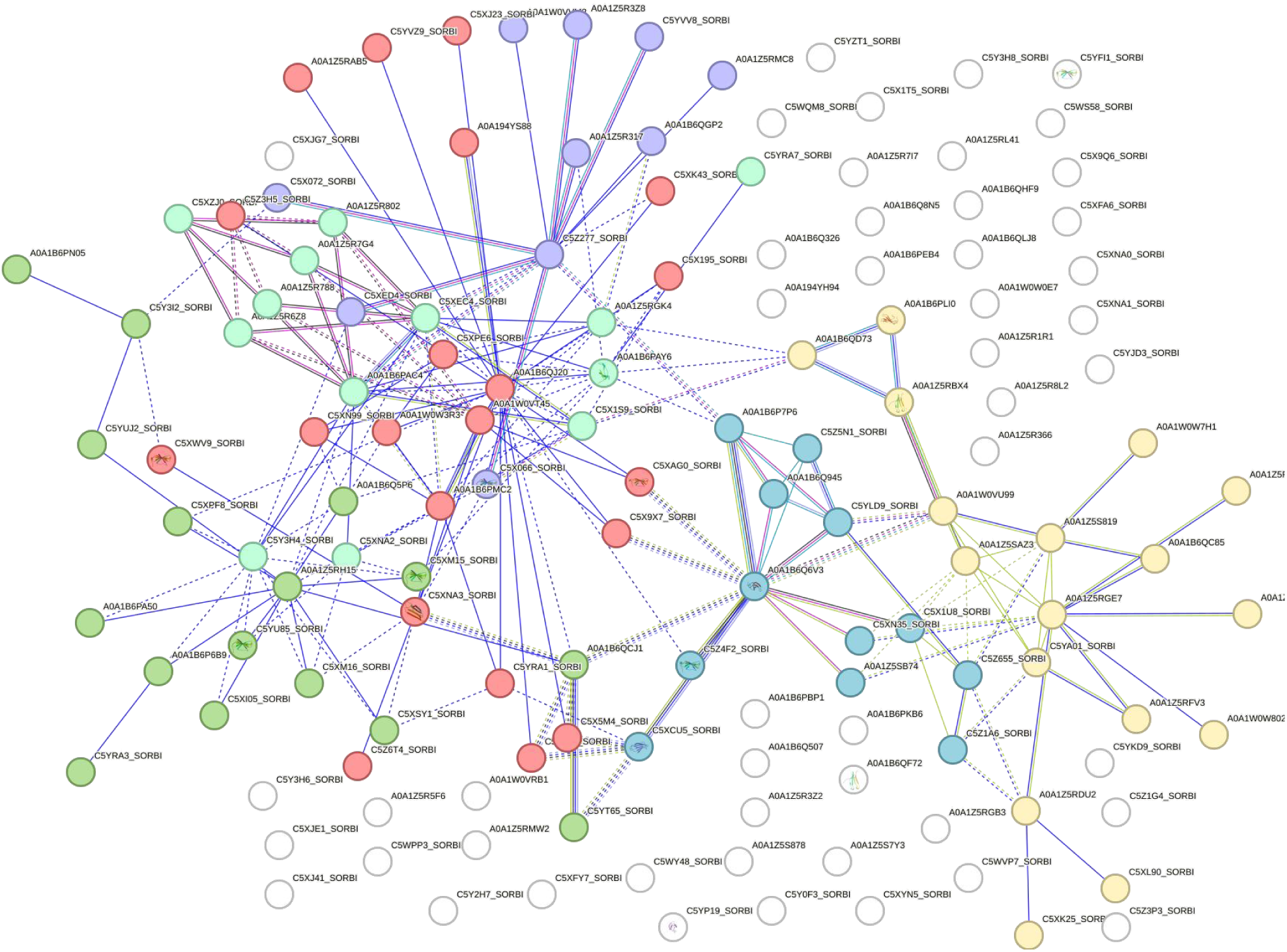
Prediction analysis of interactions between VQ and WRKY proteins in sorghum.

## Discussion

We identified 40 *SbVQ* genes in sorghum, a number comparable to Arabidopsis^[27]^, maize^[28]^ and rice^[29]^, suggesting a conserved core repertoire. However, the absence of synteny with Arabidopsis indicates that the VQ family underwent independent expansion and significant divergence during the monocot-dicot split. This expansion in sorghum was primarily driven by segmental duplication, likely underpinning the functional specialization of *VQ* genes in monocot crops.

The SbVQ proteins exhibit substantial structural and physicochemical diversity despite sharing the signature FxxhVQxhTG motif. This divergence is corroborated by gene structure analysis, where most *SbVQ* genes lack introns-a feature linked to rapid evolution. Predicted tertiary structures suggest that while the conserved VQ motif forms an α-helix for protein-protein interactions, variable regions may dictate specific subcellular localization. Notably, the high instability and aliphatic indices of certain members (e.g., SbVQ12 and SbVQ28) suggest their potential roles as transient regulators in signaling cascades^[30-31]^.

Promoter analysis revealed a high prevalence of ABRE (ABA-responsive) and TGACG/CGTCA (MeJA-responsive) elements, suggesting the SbVQ family integrates hormonal signaling under stress^[32-33]^. This aligns with the observed PEG-induced expression of several *SbVQ* genes. Notably, the significant upregulation of *SbVQ37* in roots under PEG treatment implies its positive regulatory role in the drought stress. Expression profiling under salt stress revealed functional specialization among SbVQ members, particularly in the contrasting patterns between salt-tolerant (M-81E) and salt-sensitive (Roma) lines. The significantly higher expression of *SbVQ40* and *SbVQ21* in the tolerant line, along with its upregulation in leaves under severe salt stress, suggests its role as a positive regulator of salt tolerance^[34]^. Similarly, the salt-induced expression of *SbVQ34* in the endosperm implies a potential function in protecting reproductive development under stress, which is crucial for crop yield stability.

VQ proteins primarily function by interacting with transcription factors, particularly WRKYs. Our interaction prediction identified a core network involving SbWRKY42, SbWRKY79, and SbWRKY66. The conserved VQ motif forms an α-helix ideal for docking into the WRKY domain^[35-37]^. We propose that under salt stress, specific SbVQ proteins (e.g., SbVQ10/39) act as molecular switches, modulating the transcriptional activity of partner WRKYs to activate downstream stress-responsive genes involved in osmoprotectant synthesis or ROS scavenging^[38]^.

In summary, this study provides a systematic overview of the sorghum VQ family and highlights SbVQ10/34/39 as key candidates for salt tolerance. Future work should focus on the biochemical characterization of the SbVQ-SbWRKY complex and generating transgenic lines to validate its function, paving the way for engineering saline-tolerant crops.

## Disclosure statement

No potential conflict of interest was reported by the author(s).

## Declaration of funding

The work presented here was supported by the Horizontal Project:Sereening of Salt-tolerant Germplasm and Research on Supporting Cultivation Techniques inSaline-alkali Land along Expressways (2025-CXFZJT-CXJSKF-GCCB-004).

## Data availability

All data are available in the main text.

---

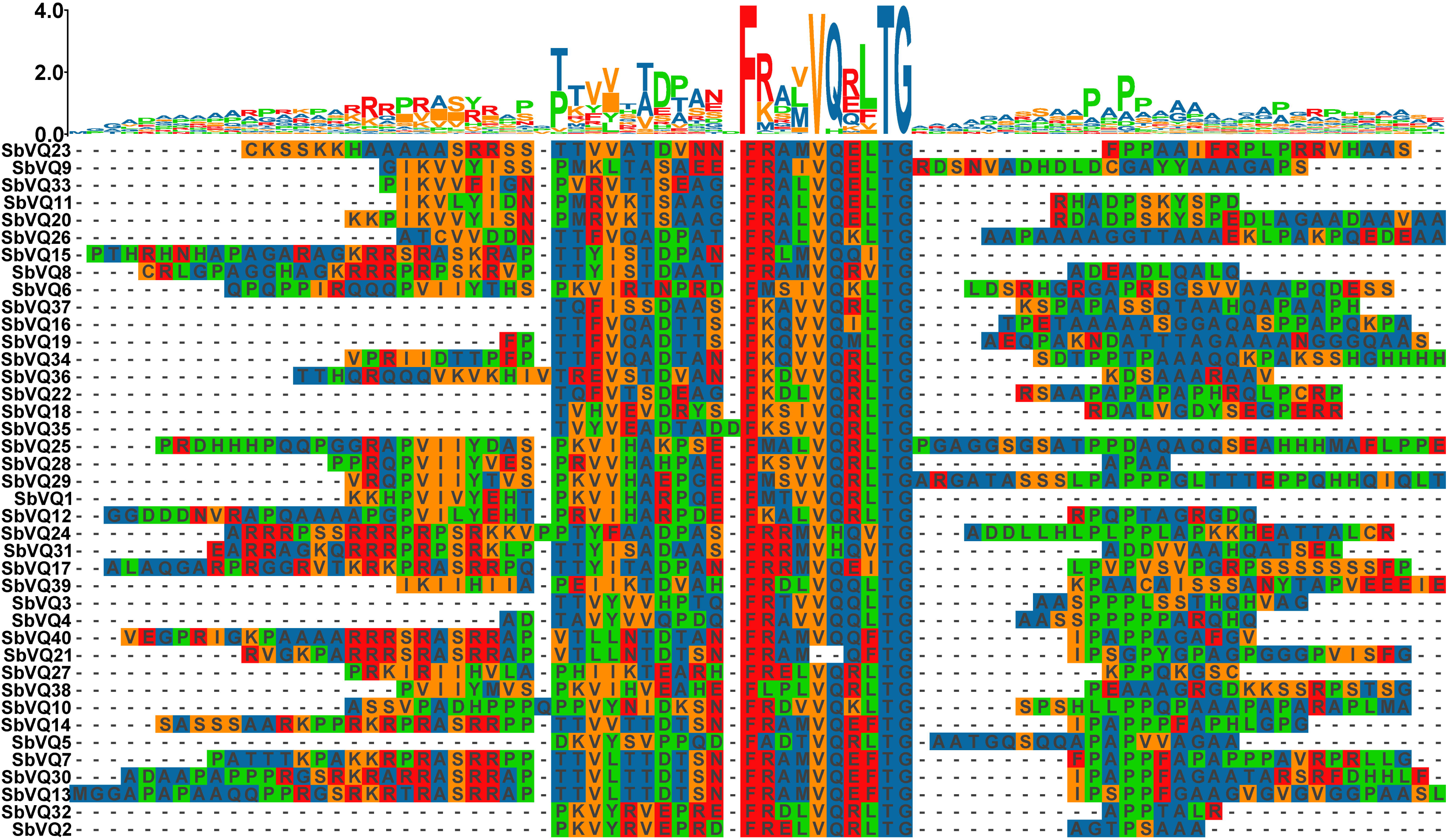

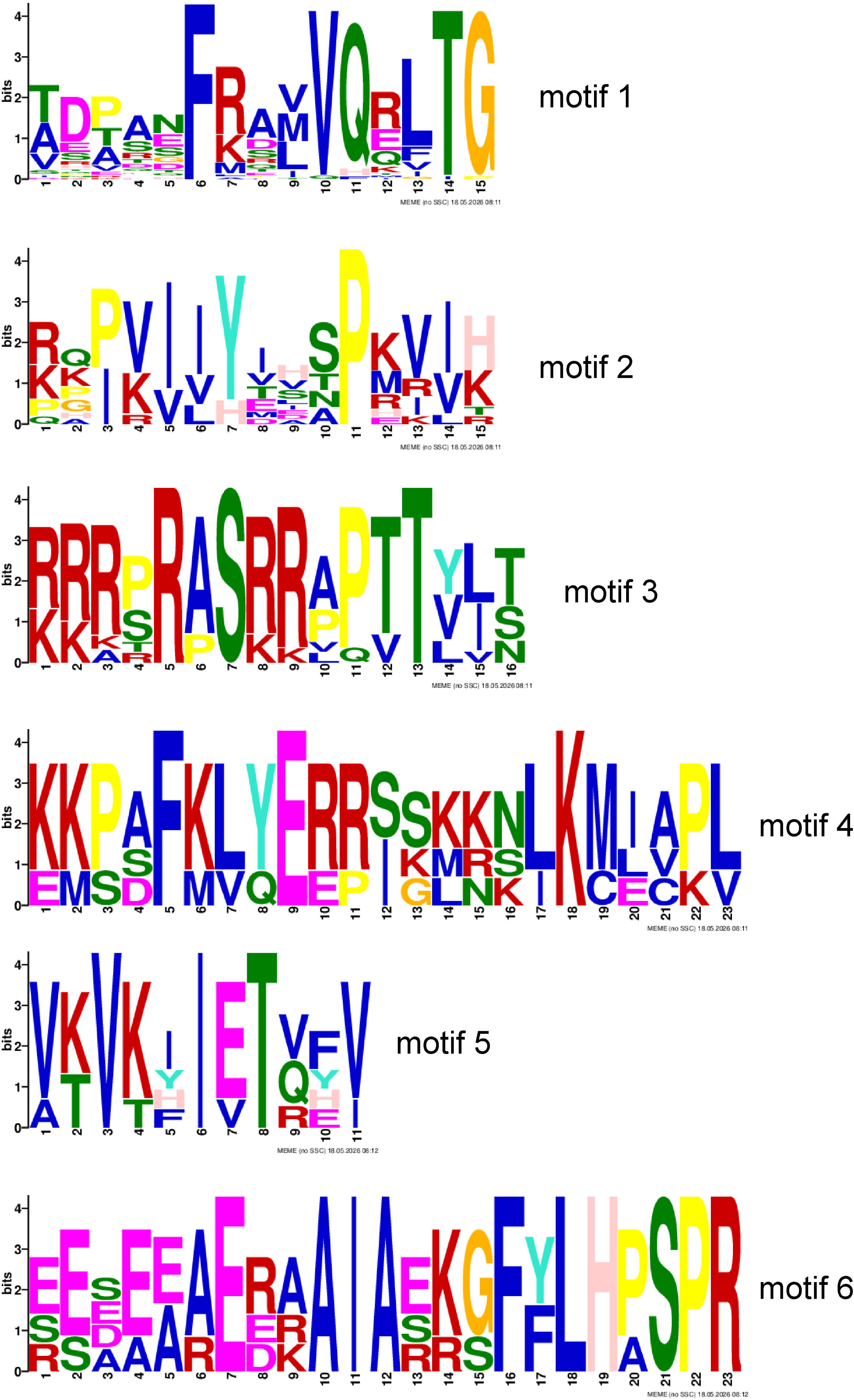

## Notes

### Competing Interest Statement

The authors have declared no competing interest.

